# Mapping a genome-scale in vivo knockout screen to a mechanistic network model identifies VAV2, RASA1, and LEPR as regulators of cardiomyocyte hypertrophy

**DOI:** 10.64898/2026.09.24.754086

**Authors:** Lionel D. Watkins, Jeffrey J. Saucerman

**Affiliations:** Department of Biomedical Engineering, University of Virginia; Division of Cardiovascular Medicine, University of Virginia

**Keywords:** Cardiac hypertrophy, Network modeling, Phenotypic screening, Systems biology, Cardiomyocytes

## Abstract

Cardiomyocyte hypertrophy is a leading clinical predictor of heart failure, yet newly identified candidate genes often remain disconnected from the signaling mechanisms that govern cardiomyocyte growth. We developed a computational-experimental pipeline that integrates genome-scale mouse knockout phenotypes with a logic-based differential equation model of hypertrophic signaling. Among 9,605 genes evaluated by the International Mouse Phenotyping Consortium, 939 knockout lines induced abnormal heart morphology. Directional curation of hypertrophy-related sub-phenotypes followed by interaction-based network expansion mapped 37 genes to the signaling model. Virtual knockdown screening identified five candidates with concordant in vivo and in silico effects: LRIG1 and CBL as predicted negative regulators and VAV2, RASA1, and LEPR as predicted positive regulators. Mechanistic subnetwork analysis linked these candidates to distinct receptor-proximal, Ras, PI3K-AKT, and MAPK signaling axes. In neonatal rat cardiomyocytes, siRNA-mediated depletion of VAV2, RASA1, or LEPR reduced phenylephrine-induced cell growth, supporting their cell-autonomous contribution to hypertrophy. Quantitative phenotyping further validated the predicted decreased cardiac hypertrophy for VAV2 and LEPR knockouts but identified potential age-dependent mechanisms for RASA1 knockout. Overall, this study establishes the application of network models to translate from in vivo phenotypic screens into pathway mechanisms.

## INTRODUCTION

Cardiac hypertrophy is an increase in myocardial mass that can initially compensate for physiological or pathological stress, but it can progress to maladaptive remodeling, contractile dysfunction, and heart failure [1–4]. Pathological hypertrophy is driven by mechanical load, neurohormonal stimulation, redox signaling, and genetic perturbations that activate interconnected signaling pathways [2,4–12]. Current therapies, including beta-blockers and inhibitors of the renin-angiotensin system, reduce hemodynamic and neurohormonal stress but do not directly resolve all intracellular mechanisms that sustain pathological cardiomyocyte growth [2,12].

Cardiomyocyte hypertrophy is governed by an interconnected network that includes G-protein-coupled receptors, Ras-MAPK, PI3K-AKT, calcium-calcineurin-NFAT, redox, and cytoskeletal signaling [2,4–12]. These pathways exhibit extensive crosstalk, feedback, and context dependence, making it difficult to infer a gene’s phenotypic effect from a single interaction. Logic-based differential equation models provide a practical way to integrate curated signaling knowledge, simulate network-wide perturbations, and generate context-specific mechanistic predictions [13–24].

The International Mouse Phenotyping Consortium (IMPC) is systematically generating and phenotyping global knockout mouse lines at a genome-scale, creating an extensive resource for connecting mammalian genes to organism-level traits and human disease [25–36]. However, individual phenotype annotations do not necessarily reveal the responsible cell type or molecular pathway. Literature-curated interaction resources and pathway-reconstruction tools can help bridge this gap by linking phenotype-associated genes to established signaling networks [37–42].

Here, we developed a computational-experimental pipeline that integrates IMPC knockout phenotypes with a published logic-based model of cardiomyocyte hypertrophy. Candidate genes were connected to the model through directed signaling interactions, screened by virtual knockdown, and prioritized by directional agreement between the simulated response and the mouse phenotype. Mechanistic subnetworks were then generated for the highest-priority candidates. Finally, the predicted positive regulators VAV2, RASA1, and LEPR were tested by siRNA knockdown in phenylephrine-stimulated neonatal rat cardiomyocytes, followed by qPCR and high-content measurement of cell area. This approach uses independent in vivo, in silico, and cell-based evidence to prioritize genes and propose testable signaling mechanisms.

## METHODS

### Processing of IMPC knockout screen

In vivo phenotypic data were obtained from the International Mouse Phenotyping Consortium (IMPC) [25–36]. We queried global knockout mouse lines associated with the primary phenotype “abnormal heart morphology” and recorded available cardiovascular subphenotypes related to heart size or shape. These included ventricular wall thickness, heart weight, chamber dimensions, and other morphology-relevant traits. Each statistically significant subphenotype was manually classified according to whether its direction was consistent with increased hypertrophy, decreased hypertrophy, or neither. Genes with at least one directionally interpretable phenotype were designated “IMPC hypertrophy genes” for comparison with network-model predictions.

To complement the initial qualitative interpretations provided by IMPC, we extracted heart-weight measurements for RASA1-KO heterozygotes at 17-18 weeks and LEPR-KO homozygotes at 11-12 weeks from the raw IMPC data. We also extracted echocardiographic measurements of diastolic left ventricular anterior wall thickness (LVAWd) and diastolic left ventricular posterior wall thickness (LVPWd) for VAV2-KO homozygotes at 9-10 weeks from the IMPC data portal.

### Network Model Incorporation

To predict pathways through which IMPC hypertrophy genes could influence cardiomyocyte growth, we mapped them to a published logic-based differential equation model of hypertrophic signaling [13–24]. Directed activating and inhibitory interactions were obtained from OmniPath [37,38]. The PathLinker algorithm, implemented in Cytoscape, was used to identify directed paths from IMPC genes (source nodes) to genes already represented in the hypertrophy model (target nodes), prioritizing direct, high-confidence connections [39–42]. For each mapped gene, the corresponding node and PathLinker-predicted reaction were added to a separate copy of the parent model in Netflux.

### Knockdown Simulations and Determination of Candidate Regulators

Knockdown simulations were performed in Netflux using a normalized-Hill logic-based ordinary differential equation framework [15]. Each IMPC-expanded model was simulated independently. The added IMPC gene was assigned a baseline input weight, and the model was simulated to steady state to obtain baseline activities for all network species.

After simulating to an initial steady state, the maximal activity parameter for the given IMPC gene was set to zero, and the simulation was continued to the new steady state. The “Cell Area” node was used as the primary metric of hypertrophy, as it is widely used in vitro and in vivo. Genes predicted to induce a >2-fold increase in cell area upon knockdown were classified as negative regulators of hypertrophy, whereas genes predicted to induce a >2-fold decrease in cell area upon knockdown were classified as positive regulators of hypertrophy. Network predictions were compared to IMPC knockout in vivo phenotypes as well as subsequent validations in cultured cardiomyocytes.

### Network visualization

Network-wide visualization of candidate regulator effects was performed in Cytoscape. For each candidate regulator, an input weight was selected by identifying the value that produced at least 90% of the maximal change in cell area across the full range of input weights, effectively establishing an EC90 input weight for visualization. Using this input, steady-state activity changes resulting from candidate regulator knockdown were mapped onto the network, and nodes were colored according to the magnitude and direction of their activity change to illustrate network-wide hypertrophic effects.

To define the pathways through which each candidate regulator influenced hypertrophy, mechanistic subnetworks were generated by integrating two complementary simulation outputs. First, we identified network nodes whose steady-state activity changed following candidate-gene knockdown. Second, each network species was individually knocked down to determine whether it altered the candidate-induced change in cell area. Nodes that both responded to candidate knockdown and contributed to propagation of the cell-area response were retained with their associated edges. All simulation and network-visualization code is available under an open-source MIT license at https://github.com/saucermanlab/IMPCnetwork.

### Neonatal rat cardiomyocyte experiments

All animal care and experiments were conducted in accordance with University of Virginia Animal Care and Use Committee policy under an approved animal protocol. Neonatal rat cardiomyocytes (NRCMs) were isolated from 1-to 2-day-old Sprague-Dawley rats using the NeoMyocyte Isolation Kit (Cellutron). Cells were seeded at 30,000 cells per well in 96-well plates in growth medium containing 15% serum.

After 24 h, dead cells and debris were removed by media exchange. Primary NRCMs are widely used to quantify stimulus-induced hypertrophic growth and signaling responses [43,46].

### SiRNA Experiments

Gene knockdown was performed using Silencer Select siRNAs (ThermoFisher), with two independent siRNA sequences used per target gene. Prior to transfection, NRCMs were serum starved for 16 h in serum-free WEB medium (Williams’ E supplemented with B-supplement) and maintained in this medium for the duration of the experiment.

For siRNA treatment, cells were transfected with 20 nM siRNA using 0.15 µL Lipofectamine RNAiMAX per well in a 96-well plate. Hypertrophic stimulation was induced concurrently with 10 µM phenylephrine. Cells were incubated under these conditions for 48 h. At the end of treatment, cells were fixed with 4% paraformaldehyde for downstream immunofluorescence imaging. Control conditions included untreated cells (no Lipofectamine, no siRNA), BLOCK-iT Fluorescent Negative Control siRNA, and Silencer Select Negative Control #1. All siRNA treatments and controls were performed in five replicates.

### Immunofluorescence and microscopy

NRCMs were fixed in 4% paraformaldehyde for 20 min at room temperature. Cells were then permeabilized with 0.1% Triton X-100 for 15 min. Cells were blocked for 1 hour in 1% bovine serum albumin (BSA) in PBS prior to primary antibody incubation. Primary antibody (α-Actinin) was probed using a goat anti-mouse primary antibody diluted 1:200, applied overnight at 4 °C. After primary incubation and washes, cells were blocked again in 5% goat serum for 1 hour and then incubated with Alexa Fluor 488- or 568-conjugated goat anti-mouse secondary antibodies (1:200) for 1 hour at room temperature. Following additional washes, nuclei were stained with DAPI for 10 mins. Plates were washed and stored in PBS until imaging. High-content imaging was performed using a PerkinElmer Operetta with a 10x objective. Images were acquired from the central region of each well in brightfield, Alexa Fluor 488 or 568, and DAPI channels.

### High-content image analysis

Images were imported in TIFF format into CellProfiler for automated image analysis [44]. Cardiomyocytes were identified by primary segmentation of nuclei and secondary segmentation of cell boundaries using cytoplasmic alpha-actinin signal. Individual-cell measurements were aggregated as the median cell area within each well, and each independently treated well was treated as the experimental unit. Downstream analyses were performed using custom Python scripts. Automated high-content measurement of cardiomyocyte area has previously been used to distinguish signaling-dependent hypertrophic phenotypes [45].

### qPCR

Total RNA was isolated from NRCMs using the Qiagen RNeasy Plus Mini Kit according to the manufacturer’s protocol. RNA quantity and purity were assessed via NanoDrop spectrophotometry, and samples were stored at −80 °C until further analysis. cDNA was synthesized from purified RNA using the iScript™ Reverse Transcription Supermix (Bio-Rad). Quantitative PCR was performed on an Agilent real-time PCR system using gene-specific primer sets. Ct values were obtained for each reaction, with β-actin serving as the housekeeping control gene. Relative gene expression was calculated using the ΔΔCt method, normalizing each target gene to β-actin and comparing siRNA-treated samples to the corresponding negative control siRNA condition.

### Statistical analysis

Statistical analyses were performed for both hypertrophy and qPCR experiments using custom Python scripts. For hypertrophy experiments, statistical significance was assessed by comparing each gene-specific siRNA treatment to its corresponding negative control condition. Each treatment was measured across two independent cell isolations with 5 wells per isolation (n = 10 total replicates per condition). A two-way ANOVA was used to account for variation across biological replicates and isolations while comparing experimental conditions, followed by Dunnett’s post hoc test to correct for multiple comparisons. A significance threshold of P < 0.05 was applied for all analyses.

For qPCR experiments, each gene-targeting siRNA was compared with the corresponding negative-control siRNA using a one-tailed Welch’s t-test. Statistical significance was defined as P < 0.05. Error bars represent standard deviations unless otherwise stated. All statistical analyses were conducted using Python-based workflows. Quantitative mouse phenotypes in **Figure 6B and C** were compared using two-tailed t-tests with Sidak correction for multiple comparisons.

## RESULTS

### IMPC phenotype curation identifies genes associated with altered cardiac morphology

To identify candidate regulators of cardiac hypertrophy from large-scale in vivo phenotyping, we queried the IMPC database for murine global knockout lines annotated with abnormal heart morphology. We processed the 9,605 genes represented in the dataset for knockouts associated with “abnormal heart morphology”. Because “abnormal heart morphology” includes findings that are not necessarily caused by cardiac growth, we manually evaluated related available cardiovascular sub-phenotypes, including ventricular wall thickness, heart weight, and chamber dimensions. Genes with at least one statistically significant, directionally interpretable hypertrophy-related sub-phenotype were retained as IMPC hypertrophy genes (939 genes). For each retained gene, loss of function was classified as being associated with increased or decreased hypertrophy. This curation converted heterogeneous organism-level annotations into directional phenotypes suitable for comparison with network simulations (**Figure 1**).

**Figure 1.**
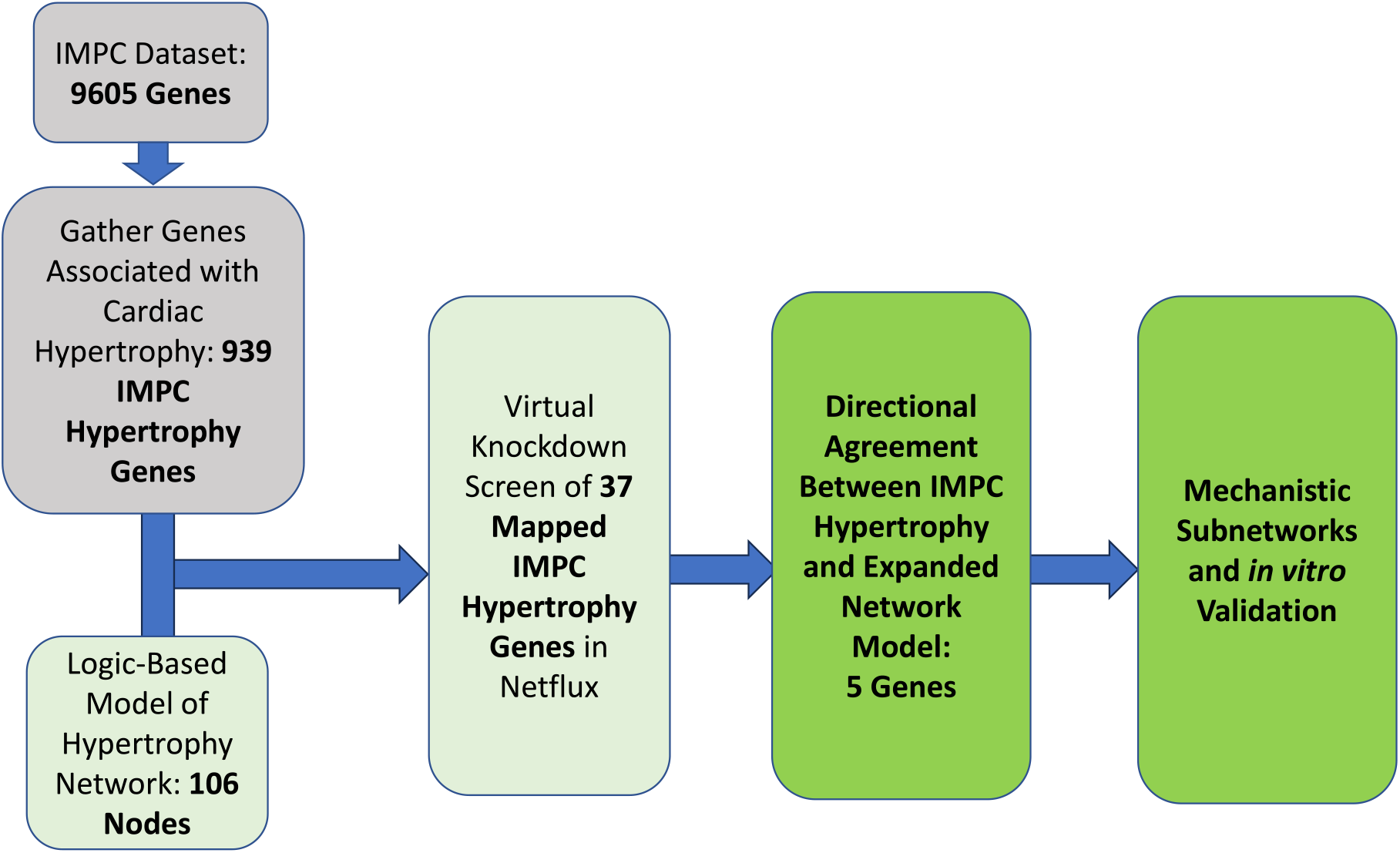
Integration of IMPC phenotypic data with a logic-based model identifies candidate regulators of cardiomyocyte hypertrophy. Schematic of the study workflow. The International Mouse Phenotyping Consortium (IMPC) contained in vivo knockout data for several thousand genes, a subset of which were able to be curated as IMPC hypertrophy genes. After mapping these genes onto a validated model of cardiomyocyte hypertrophy, further analysis allowed for identification of novel candidate regulators, for which mechanistic subnetwork analysis was performed.

### Network expansion maps 37 IMPC hypertrophy genes to hypertrophy signaling

The curated IMPC gene set did not by itself explain how individual genes might regulate cardiac growth. We therefore mapped the IMPC hypertrophy genes to a previously published logic-based differential equation model of cardiomyocyte hypertrophy [13]. The parent model contains 106 signaling nodes corresponding to 212 genes and representing proteins, genes, small molecules, and phenotypic outputs. It integrates GPCR, Ras-MAPK, PI3K-AKT, calcium-dependent, and transcriptional pathways that converge on cell area.

Directed signaling interactions from OmniPath were searched with PathLinker to connect IMPC genes to nodes already represented in the hypertrophy model [37–40]. Prioritizing direct connections identified 37 IMPC genes with first-neighbor links to the parent network. (**Figure 2A**). Each mapped gene and its connecting reaction were added separately in Netflux, generating 37 gene-specific expanded models. This design retained a common parent network while allowing the effect of each newly mapped gene to be evaluated independently.

**Figure 2.**
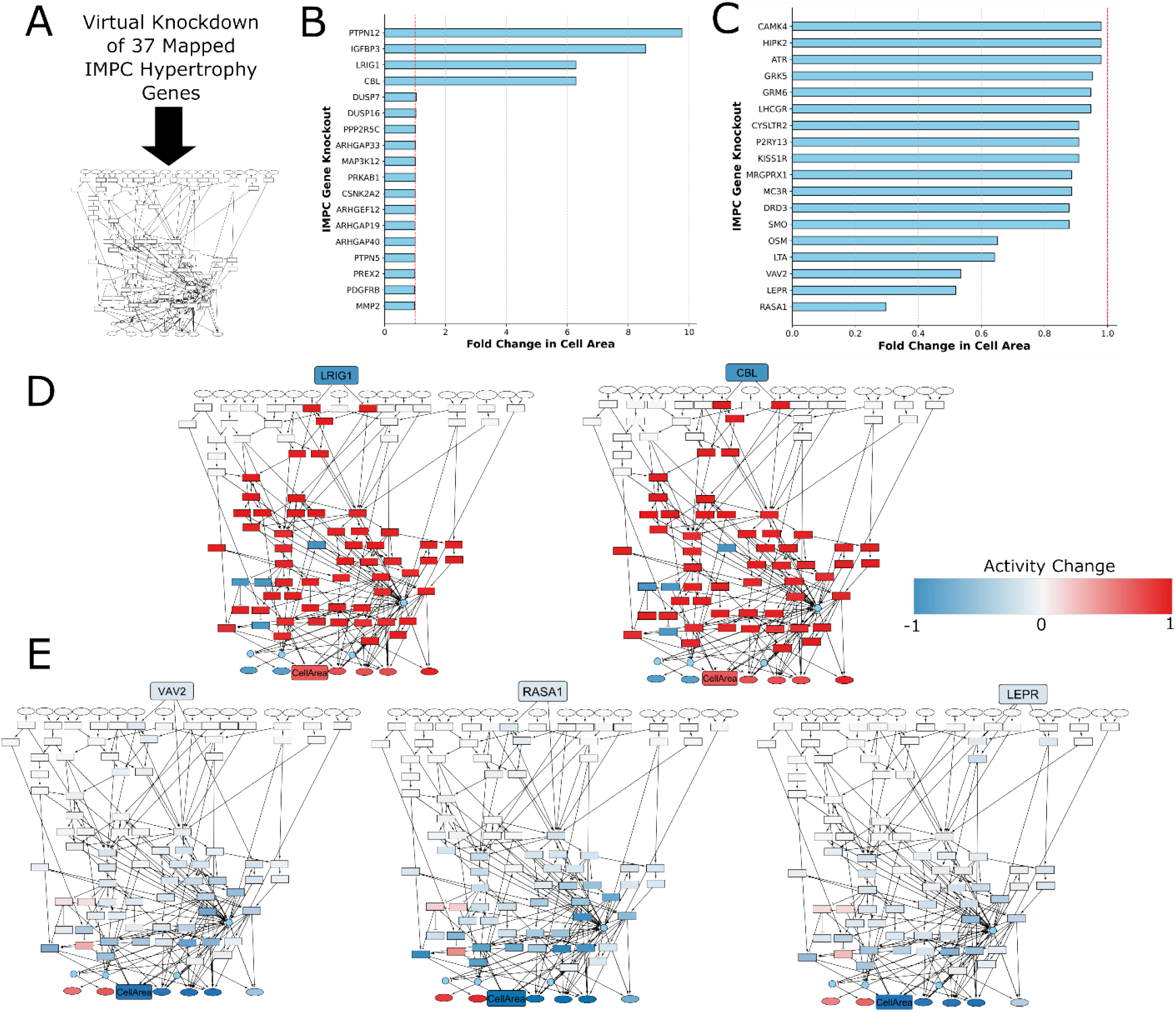
Virtual knockdown screening of IMPC genes predicts network-wide hypertrophic effects to identify positive and negative regulators of cardiomyocyte hypertrophy. (A) 37 mapped IMPC hypertrophy genes were individually knocked down in the logic-based cardiomyocyte hypertrophy network. Simulated knockdown results are shown for (B) negative regulators of hypertrophy and (C) positive regulators of hypertrophy. Network-wide activity changes are visualized for (D) the negative regulators LRIG1 and CBL and (E) the positive regulators VAV2, LEPR, and RASA1.

### Virtual knockdown screening identifies five candidate regulators with concordant in vivo and in silico effects

For each of the 37 expanded models, the added IMPC hypertrophy gene was assigned a baseline input reaction weight and the model was simulated to steady state. The gene was then computationally knocked down by setting its maximal activity to zero, and the model was simulated to a new steady state. The hypertrophic response was quantified as the fold change in cell area relative to baseline. A decrease in cell area following knockdown indicated that the gene functioned as a predicted positive regulator of hypertrophy, whereas an increase indicated a predicted negative regulator.

Virtual knockdown responses varied substantially across the 37 mapped genes (**Figure 2B,C**). Knockdown of predicted positive regulators decreased cell area, whereas knockdown of predicted negative regulators increased cell area. These directionally distinct responses provided the basis for comparison with the corresponding IMPC phenotypes.

Five genes showed concordant response directions in the in vivo and in silico datasets: LRIG1, CBL, VAV2, RASA1, and LEPR. VAV2, RASA1, and LEPR were classified as positive-regulator candidates because their loss reduced hypertrophy in both datasets. LRIG1 and CBL were classified as negative-regulator candidates because their loss increased hypertrophy. Network-wide visualization showed that each candidate altered a distinct pattern of signaling activity, indicating that similar effects on cell area could arise through different combinations of intermediate nodes (**Figure 2D,E**).

### Mechanistic subnetworks predict distinct routes from candidate regulators to cardiomyocyte growth

To distinguish network responses correlated with candidate-gene knockdown from nodes required to transmit its effect on cell area, we performed mechanistic subnetwork analysis. For each candidate, we first identified nodes whose steady-state activity changed following candidate knockdown. We then independently knocked down each network node and measured the effect of that perturbation on the candidate-induced change in cell area. The intersection of responsive nodes and nodes that regulated the candidate phenotype defined the mechanistic subnetwork (**Figure 3A**).

**Figure 3.**
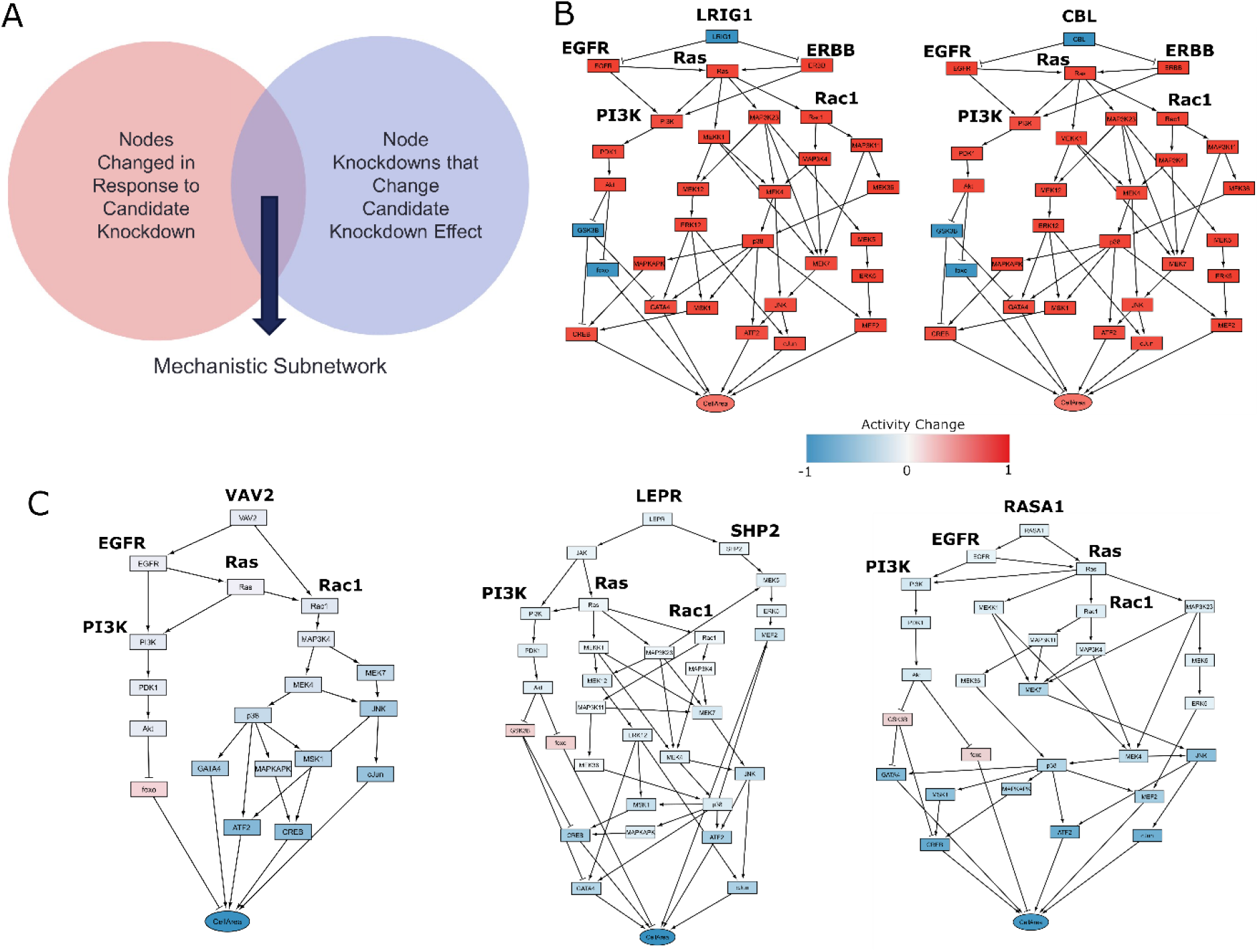
Mechanistic subnetwork analysis identifies the pathways that mediate the hypertrophic effects of candidate genes. (A) Mechanistic subnetworks were defined as the intersection of nodes that change in response to candidate gene knockdown and nodes whose individual knockdown affect the candidate gene knockdown effect on cell area. Individual mechanistic subnetworks are shown for B) negative regulators (LRIG1, CBL) and C) positive regulators (VAV2, LEPR, RASA1). Directed edges represent either activating interactions (arrows) or inhibiting interactions (blunted).

The LRIG1 and CBL subnetworks converged on receptor-proximal EGFR/ERBB signaling and propagated through Ras, PI3K-AKT, and MAPK modules to transcriptional regulators and cell area (Figure 3B). Although the two subnetworks were similar, their placement upstream of receptor signaling suggests that LRIG1 and CBL may regulate the strength or duration of common growth-factor inputs. The VAV2 subnetwork connected receptor-proximal signals to Ras and Rac1 and then to MAPK branches, including MAP3K4, p38, JNK, and downstream transcription factors. The LEPR subnetwork engaged JAK- and SHP2-associated routes together with PI3K-AKT and Ras-MAPK signaling, whereas the RASA1 subnetwork was organized around EGFR/Ras with branches through PI3K-AKT and multiple MAPK modules (**Figure 3C**). These subnetworks constitute model-derived hypotheses about experimentally testable pathway dependencies rather than evidence of direct biochemical causality.

### siRNA knockdown verifies reduced expression of positive-regulator candidates

We next tested the three positive-regulator candidates in neonatal rat cardiomyocytes (NRCMs). Cells isolated from 1-to 2-day-old Sprague-Dawley rats were plated in 96-well format, serum-starved, and treated concurrently with 20 nM siRNA and 10 µM phenylephrine for 48hr (**Figure 4A**). Two independent siRNA sequences were evaluated for each candidate. In parallel cultures, target-gene expression was measured by qPCR and normalized to beta-actin using the delta-delta Ct method.

**Figure 4.**
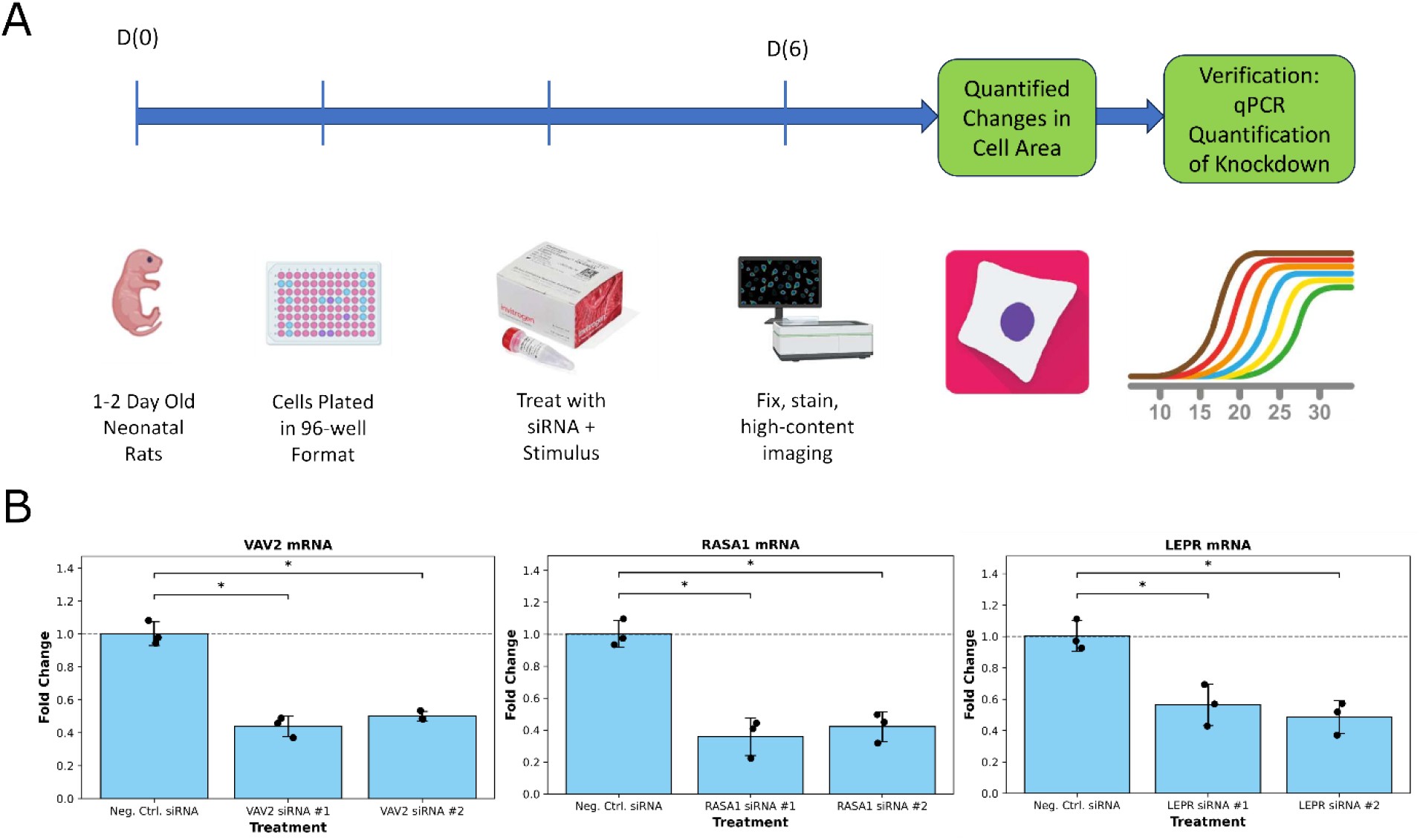
siRNA-mediated knockdown reduces VAV2, RASA1, and LEPR transcript expression in neonatal rat cardiomyocytes. (A) Experimental workflow. Cardiomyocytes isolated from 1- to 2-day-old neonatal rats were plated in 96-well plates, treated with gene-targeting siRNA and phenylephrine, fixed and stained for high-content imaging, and analyzed for cell area. Target-gene knockdown was evaluated by qPCR. (B) Relative VAV2, RASA1, and LEPR mRNA expression following treatment with negative-control siRNA or two independent gene-targeting siRNAs. Expression was normalized to beta-actin and to the corresponding negative-control condition, represented by the dashed line at a fold change of 1. Individual points represent replicate measurements (n = 3). Bars show mean +/− SD. Comparisons were performed using a one-tailed Welch’s t-test. *P < 0.05.

Both siRNAs targeting each candidate significantly reduced the corresponding transcript relative to negative-control siRNA (P < 0.05; **Figure 4B**). Mean expression after knockdown ranged from approximately 0.36- to 0.57-fold of the negative-control condition. These results confirmed target engagement for VAV2, RASA1, and LEPR under the experimental conditions used for the hypertrophy assay.

### Knockdowns of VAV2, RASA1, and LEPR attenuate PE-induced cardiomyocyte hypertrophy

Following siRNA treatment and phenylephrine stimulation, NRCMs were stained for alpha-actinin and DAPI. High-content images were acquired with an Operetta platform, and CellProfiler was used to segment cardiomyocytes and quantify cell area [44,45]. Representative images from two independent cell isolations showed reduced cardiomyocyte size following knockdown of VAV2, RASA1, or LEPR compared with the phenylephrine-treated negative-control condition (**Figure 5A**).

**Figure 5.**
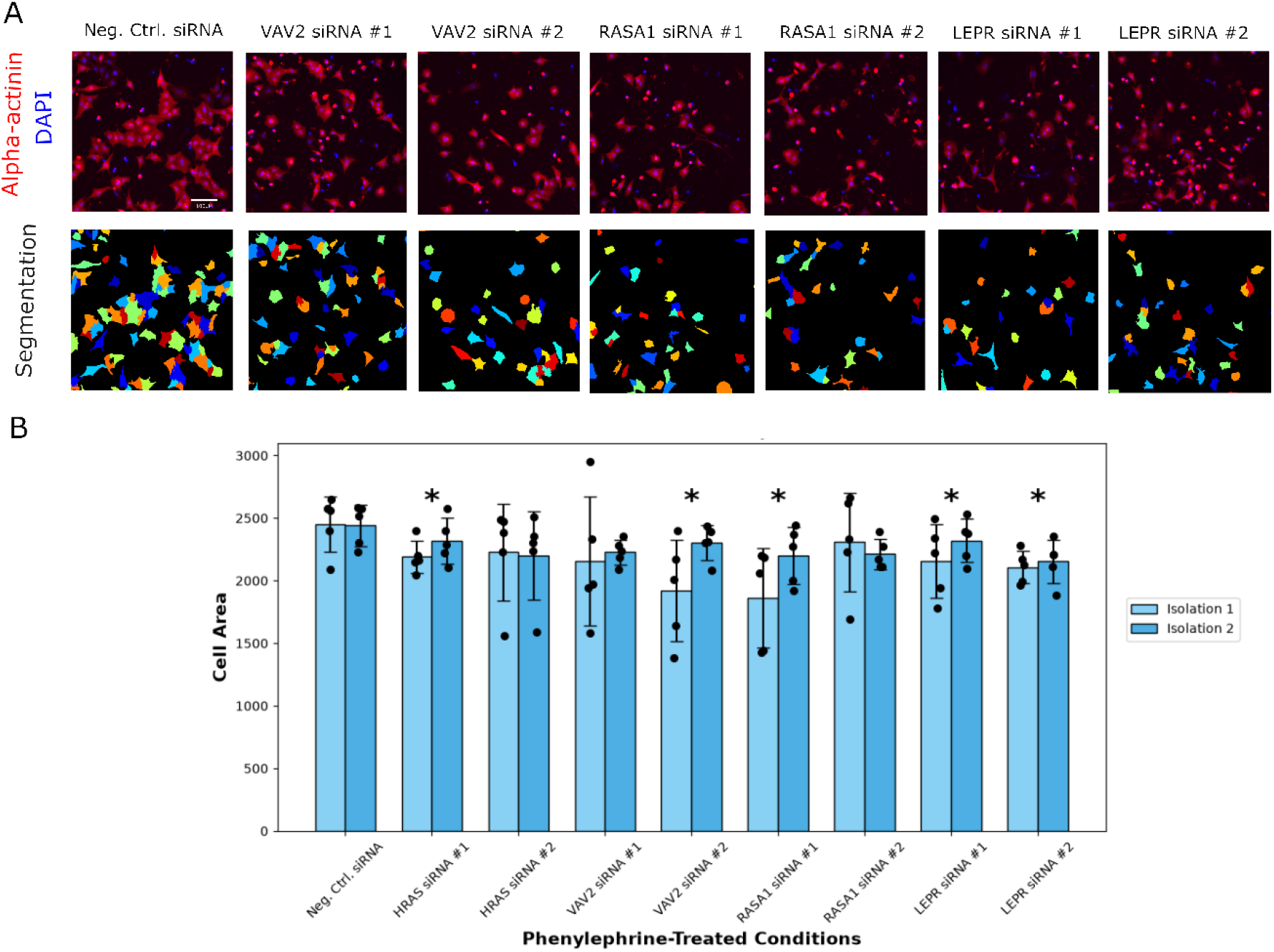
Knockdown of VAV2, RASA1, and LEPR attenuates phenylephrine-induced cardiomyocyte hypertrophy. (A) Representative fluorescence images and cell segmentation of PE-treated neonatal rat cardiomyocytes transfected with negative-control siRNA or independent siRNAs targeting VAV2, RASA1, or LEPR. Cardiomyocytes were identified by α-actinin staining (red), and nuclei were stained with DAPI (blue). Each color in the segmentation images represents an individually identified cardiomyocyte. Scale bar, 100 µm. (B) Quantification of cardiomyocyte cell area following knockdown of HRAS, VAV2, RASA1, or LEPR in two independent biological isolations. HRAS siRNA served as a positive-control perturbation. Individual points represent replicate wells (n = 5 per condition within each isolation); bars show mean ± SD. Statistical comparisons with the PE-treated negative-control siRNA condition were performed using two-way ANOVA followed by Dunnett’s post hoc test. *P < 0.05.

Five replicate wells were analyzed per condition in each of two independent cell isolations. Median cell area within each well was used as the well-level response, and data were analyzed by two-way ANOVA followed by Dunnett’s post hoc test. VAV2 siRNA #2, RASA1 siRNA #1, and both LEPR-directed siRNAs significantly reduced cell area relative to negative-control siRNA (P < 0.05; **Figure 5B**). HRAS siRNA #1 also reduced cell area, whereas HRAS siRNA #2, VAV2 siRNA #1, and RASA1 siRNA #2 did not reach the prespecified significance threshold. Together with the qPCR results, these findings support positive-regulator roles for VAV2, RASA1, and LEPR in phenylephrine-stimulated cardiomyocyte growth while also demonstrating sequence-dependent differences in phenotypic effect.

### Validation of cardiac phenotypes for VAV2, RASA1, and LEPR knockout mice

The initial IMPC classification was based on statistically significant hypertrophy-related subphenotypes rather than reanalysis of every underlying quantitative dataset. We therefore examined available quantitative cardiac measurements for VAV2-, LEPR-, and RASA1-knockout mice. Because the IMPC portal did not provide the same measurements, ages, zygosities, or sample sizes for every line, these comparisons were interpreted as gene-specific corroborative analyses rather than a uniform validation cohort.

Diastolic left ventricular anterior wall thickness (LVAWd) and posterior wall thickness (LVPWd) were evaluated in 9- to 10-week-old VAV2-knockout mice (**Figure 6A,B**). VAV2-knockout males and females exhibited significantly reduced LVAWd and LVPWd relative to wild-type mice (P < 0.05), consistent with the predicted and experimentally observed reduction in hypertrophic growth following VAV2 loss.

**Figure 6.**
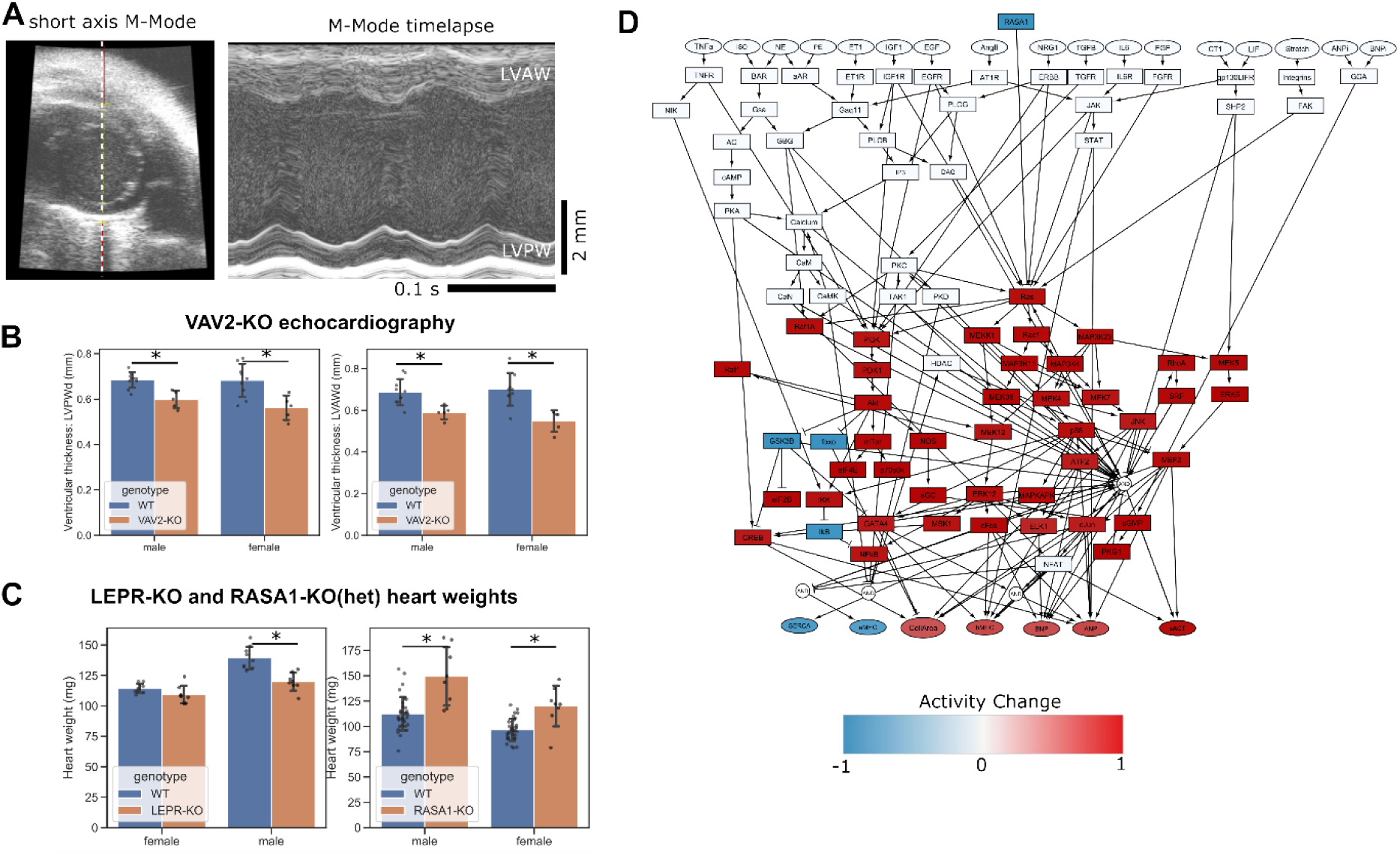
Quantitative phenotyping of VAV2, LEPR, and RASA1 knockout mice. (A) Representative short-axis M-mode echocardiographic image and M-mode time-lapse recording from a wild-type C57BL mouse. LVAW, left ventricular anterior wall; LVPW, left ventricular posterior wall. Scale bars, 2 mm and 0.1 s. (B) Quantification of diastolic left ventricular posterior wall thickness (LVPWd) and diastolic left ventricular anterior wall thickness (LVAWd) in 9- to 10-week-old wild-type mice (n = 10 male and n = 9 female) and VAV2-knockout mice (n = 5 male and n = 5 female). (C) Heart weights of wild-type and homozygous LEPR-knockout mice and of 17- to 18-week-old wild-type and heterozygous RASA1-knockout mice. (D) Revised network model that removes RASA1-EGFR interaction and makes RASA1 a negative regulator of RAS is sufficient to predict increased Ras activity and cardiomyocyte hypertrophy. Node color represents the change in modeled activity between baseline and knockdown contexts. Statistical comparisons were performed using two-tailed t-tests with Sidak correction; *P < 0.05.

Heart weight provided a complementary in vivo measure (**Figure 6C**). LEPR-knockout males had lower heart weights than wild-type males (P = 0.0005), whereas the difference in females was not significant (P = 0.134). Both male and female LEPR-knockout mice nevertheless exhibited approximately twofold greater body mass than wild-type mice, underscoring the systemic metabolic phenotype of LEPR loss and the need to interpret unnormalized heart weight cautiously.

Both model predictions and in vitro validations demonstrated that RASA1 knockdown decreases cardiomyocyte hypertrophy in those contexts. This was further supported by representative histology images of heart size and cardiomyocyte size from RASA1-KO heterozygous mice, taken at 8 weeks of age (**Supplementary Figure S1**). In contrast, the available IMPC data from these mice were taken at 17-18 weeks of age and indicated increased heart weight from both sexes (**Figure 6C**).

To explore mechanisms that may be responsible for this discordance, we re-examined protein interaction data available for RASA1 in Omnipath. The initial model (**Figure 2**) contained RASA1 → EGFR and RASA1 → HRAS derived from annotations in Omnipath v2022. However, inspection of updated annotations in Omnipath v2026 indicated that RASA1 and EGFR interact by binding rather than directional stimulation. Second, our original annotation of consensus activation from RASA1 to HRAS based on Omnipath v2022 is listed in Omnipath v2026 instead as consensus inhibition from RASA1 to HRAS. Indeed, this relationship is subtle because RASA1 stimulates HRAS GTPase activity, which terminates downstream activity of HRAS. Thus we revised the model based on the Omnipath v2026 annotations and re-simulated the effects of RASA1 knockdown. The revised model predicted that RASA1 knockdown enhances cardiomyocyte hypertrophy, which conflicts with the in vitro data (**Figure 4**) but is consistent with the adult mouse heart weight data (**Figure 6C**). Differences in age, zygosity, whole-organ physiology, and systemic or noncardiomyocyte effects may contribute to the discordance between the adult mouse heart-weight phenotype and the cell-autonomous assay.

## DISCUSSION

In this study, we developed a computational-experimental strategy that combines directionally curated IMPC cardiac phenotypes with logic-based network modeling and targeted cardiomyocyte validation experiments. From 939 genes associated with cardiac hypertrophy in vivo, mechanistic interaction-based expansion connected 37 genes to the hypertrophy model, predicting concordance between in vivo and virtual-knockdown directions prioritized five candidates. LRIG1 and CBL were predicted negative regulators as their knockdown increased cell area, whereas VAV2, RASA1, and LEPR were predicted positive regulators as their knockdown decreased cell area. siRNA validation experiments supported positive-regulator roles for VAV2, RASA1, and LEPR in phenylephrine-stimulated NRCMs. This multi-stage design is valuable because IMPC phenotypes capture integrated whole-animal biology [25–36], while logic-based models formalize pathway crosstalk and generate testable, context-dependent mechanisms [13–24].

The VAV2 and RASA1 findings illustrate both the value and limits of network-level interpretation. VAV proteins link receptor activation to Rho-family GTPases and cardiovascular signaling [47–50], consistent with the modeled VAV2 route through Ras/Rac1 and MAP3K4-p38/JNK. Only VAV2 siRNA #2 significantly reduced cell area despite transcript reduction by both sequences, suggesting differences in protein depletion, nonlinear signaling, or sequence-specific effects. RASA1 ordinarily restrains Ras-MAPK signaling [51–54], yet the original model (**Figure 2E**) and RASA1 siRNA #1 (**Figure 5B**) supported a positive-regulator role via simulations and neonatal cardiomyocyte experiments, respectively. Review of the OmniPath v2022 and Omnipath v2026 annotations showed that the modeled RASA1-to-EGFR activation was supported only as a physical, undirected interaction, while the RASA1-to-HRAS relationship had changed from consensus activation in OmniPath v2022 to consensus inhibition in Omnipath v2026. After revising these interactions, simulated RASA1 knockdown increased cell area, changing RASA1’s context from that of a positive regulator to a negative regulator (**Figure 6D**), consistent with the increased heart weight of older RASA1-heterozygous mice (**Figure 6C**). The remaining experimental divergence may reflect age, developmental or vascular contributions, network adaptation, or siRNA-specific effects. Protein confirmation, Ras-GTP measurement, and time-resolved MAPK assays are important next steps to experimentally support these findings.

LEPR provided the most direct link between metabolism and canonical hypertrophic signaling. Leptin-receptor activation engages JAK/STAT, PI3K-AKT, and Ras-MAPK pathways and has context-dependent cardiovascular effects [55–58], matching the predicted LEPR subnetwork and the reduction in cell area produced by both LEPR siRNAs. However, the lower absolute heart weight in male LEPR-knockout mice occurred despite markedly increased body mass (data not shown), and females did not show a significant heart-weight reduction. Future analyses should distinguish LEPR isoforms, define leptin exposure, and normalize cardiac mass to appropriate body-size measures. LRIG1 and CBL were not tested in NRCMs, but their predicted negative-regulator roles are consistent with literature describing LRIG1-mediated restraint of EGFR/ERBB signaling and CBL-dependent receptor ubiquitination, internalization, and downregulation [59–63]. Effects of negative regulators are often highly context-dependent and are challenging therapeutic targets. Still these candidates can be tested further by measuring receptor abundance and trafficking together with stimulus-dependent ERK/AKT kinetics and cell area.

Several limitations constrain interpretation. Network expansion depended on existing interaction knowledge [37–42], and normalized-Hill models represent semiquantitative activities rather than measured concentrations or reaction flux [13,15]. While we mapped hypertrophy genes with length 1, more genes could be mapped with longer pathlengths with a likely cost of accuracy. Experimentally, phenylephrine represents a single α1-adrenergic stimulus [4] using cell area as the principal endpoint.

The quantitative IMPC findings did not always agree with the network simulations and cell-based experiments, and developmental age may contribute to the disagreement. Comparison across age and cells vs. hearts indicates that RASA1 may transition from being a positive to a negative regulator hypertrophy. The IMPC measurements also differed in sex, zygosity, endpoint, and sample size, which may require further dedicated in vivo studies to characterize fully. Despite these limitations, convergence across phenotype curation, simulation, and cell-based perturbation supports VAV2, RASA1, and LEPR as context-dependent regulators of cardiomyocyte growth and nominates LRIG1 and CBL for further study. Finally, this study demonstrates how mechanistic network modeling can bridge a challenging gap from in vivo genetic candidates to pathway mechanisms that may provide new therapeutic candidates for disease.

## ABBREVIATIONS

ACE: Angiotensin-converting enzyme
AKT: Protein kinase B
ANOVA: Analysis of variance
ARBs: Angiotensin receptor blockers
DAPI: 4′,6-diamidino-2-phenylindole
EGFR: Epidermal growth factor receptor
ERBB: Erb-b2 receptor tyrosine kinase
ERK: Extracellular signal-regulated kinase
GPCR: G-protein-coupled receptor
GTPase: Guanosine triphosphatase
IMPC: International Mouse Phenotyping Consortium
JAK: Janus kinase
JNK: c-Jun N-terminal kinase
MAPK: Mitogen-activated protein kinase
NFAT: Nuclear factor of activated T-cells
NRCMs: Neonatal rat cardiomyocytes
PE: Phenylephrine
PI3K: Phosphoinositide 3-kinase
qPCR: Quantitative polymerase chain reaction
RasGAP: Ras GTPase-activating protein
SHP2: SH2 domain-containing protein tyrosine phosphatase-2
siRNA: Small interfering RNA
STAT: Signal transducer and activator of transcription).

## FUNDING

This study was funded by the National Institutes of Health HL162925 (grant to JJS, supplement to LDW) and HL176700 (to JJS), and T32 GM145443 (to LDW).

## ACKNOWLEDGEMENTS

We thank Dr. Yongde Bao (UVA Genome Analysis and Technology Core), Kaitlyn Wintruba, Lavie Ngo, Bryce Murillo, and Pichayathida Luanpaisanon for their advice and technical assistance.

## Supplementary Figures

**Supplementary Figure 1.**
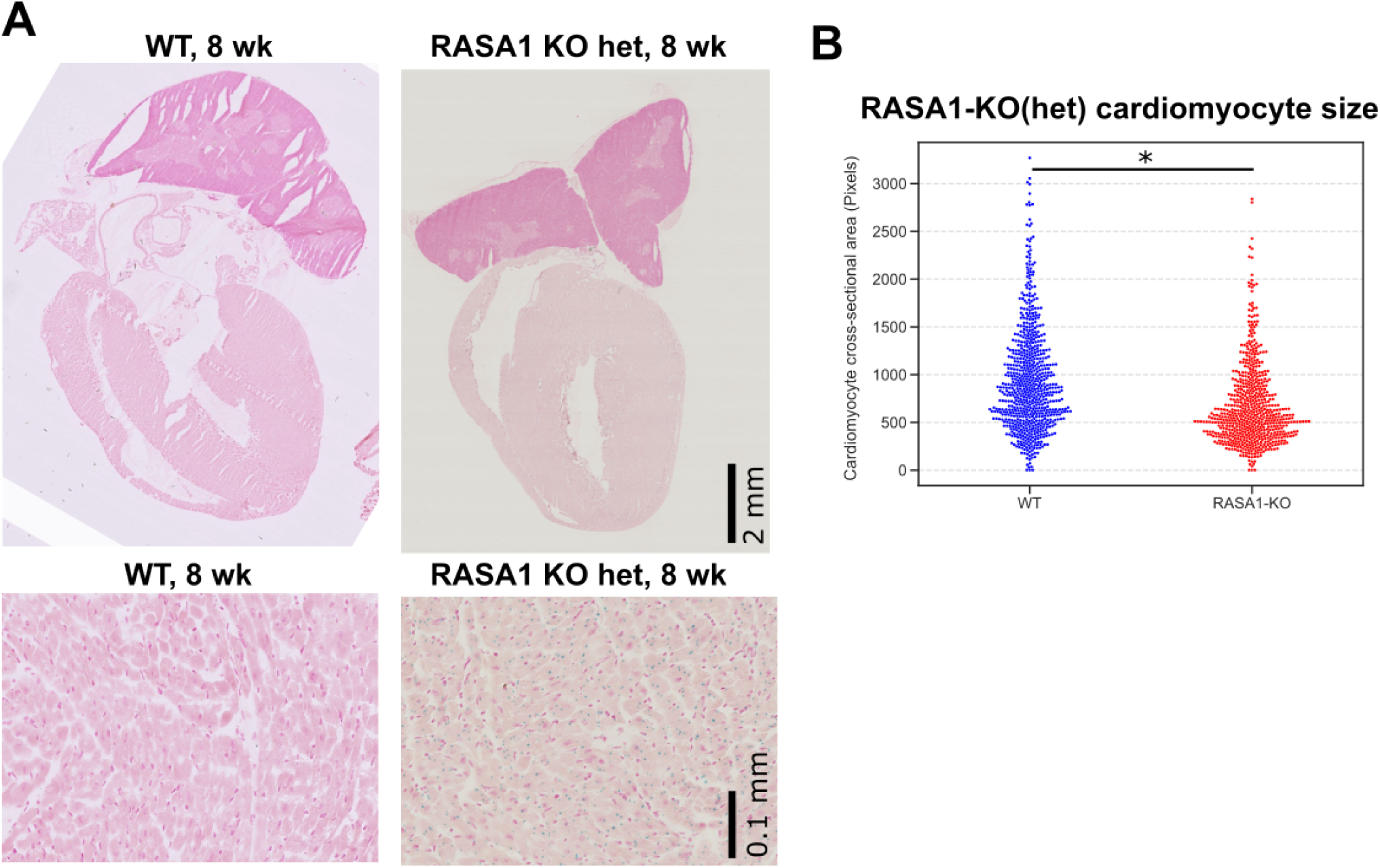
Representative tissue sections of whole hearts and cardiomyocyte cross-sections from WT and RASA1-KO heterozygous mice at 8 weeks of age. F) Quantification of cardiomyocyte cross-sectional area from Figure 6D. *p<0.05 two-tailed T test with Sidak correction, n =744 WT myocytes and n=679 RASA1-KO myocytes.

## Notes

### Competing Interest Statement

The authors have declared no competing interest.

